# An auto-release mechanism for HMCES-DNA-protein crosslinks

**DOI:** 10.1101/2022.12.18.520715

**Authors:** Maximilian Donsbach, Sophie Dürauer, Kha T. Nguyen, Florian Grünert, Denitsa Yaneva, Daniel R. Semlow, Julian Stingele

**Affiliations:** Department of Biochemistry, Ludwig-Maximilians-University Munich, 81377 Munich, Germany; Gene Center, Ludwig-Maximilians-University Munich, 81377 Munich, Germany; Division of Chemistry and Chemical Engineering, California Institute of Technology, Pasadena, 91125 CA, USA

## Abstract

The conserved protein HMCES crosslinks to abasic (AP) sites in ssDNA to prevent strand scission and the formation of toxic dsDNA breaks during replication. Here, we report a non-proteolytic release mechanism for HMCES-DNA-protein crosslinks (DPCs), which is regulated by DNA context. In ssDNA and at ssDNA-dsDNA junctions, HMCES-DPCs are stable, which efficiently protects AP sites against spontaneous incisions and cleavage by APE1 endonuclease. In contrast, HMCES-DPCs are quickly released in dsDNA, allowing APE1 to initiate downstream repair. Mechanistically, we show that release is governed by two components. First, a conserved glutamate residue within HMCES’ active site catalyses reversal of the crosslink. Second, affinity to the underlying DNA structure determines whether HMCES re-crosslinks or dissociates. Our study reveals that the protective role of HMCES-DPCs involves their controlled release upon bypass by replication forks, which restricts DPC formation to a necessary minimum.

## Introduction

Covalent crosslinks between proteins and DNA (DNA-protein crosslinks, DPCs) are dangerous lesions caused by a variety of endogenous and exogenous sources, including widely-used chemotherapeutic agents^1,2^. DPCs are toxic because they interfere with DNA replication^3^. Therefore, cells possess conserved repair mechanisms that target DPCs in replication-dependent and -independent manners^4^. DPC repair involves the proteolytic destruction of the protein adduct by DPC-specific proteases of the SPRTN/Wss1 family or by proteasomal degradation^5–8^. Failure to degrade DPCs has drastic consequences; complete loss of SPRTN is lethal in mammalian cells, while partial loss-of-function results in premature aging and predisposition to liver cancer^9–11^. Despite the severe phenotypes associated with the absence of SPRTN alone, several additional proteases appear to target DPCs^12–16^. The diversity of repair mechanisms underlines the threat posed by DPCs. However, some DPCs have important physiological roles. The human protein HMCES forms crosslinks with abasic (AP) sites to protect genome integrity^17^.

AP sites are frequent endogenous DNA lesions, which arise spontaneously or enzymatically during base excision repair and active DNA demethylation^18^. AP sites exist in equilibrium between a closed-ring furanose and an open-ring aldehyde form. The latter is prone to undergo spontaneous β-elimination resulting in strand scission and DNA single-strand break (SSB) formation, which can also occur enzymatically upon AP site cleavage by AP endonucleases/lyases^19,20^. If such SSBs form in double-stranded DNA (dsDNA), they are swiftly repaired by the cellular SSB repair machinery^21^. In contrast, incision of AP sites in ssDNA, e.g. at the replication fork, will result in the formation of toxic DSBs^22,23^. To prevent such a catastrophic scenario, the conserved catalytic SRAP (SOS response associated peptidase) domain of HMCES (Figure 1A) associates with replication forks to crosslink to AP sites in ssDNA^17^. Crosslinking occurs between the N-terminal cysteine residue of the SRAP domain (methionine is proteolytically removed) and an AP site, resulting in the formation of a thiazolidine ring, which prohibits strand scission (Figure 1B, 1C and refs^24–26^). DPC formation has been suggested to be initiated by the N-terminal amino group attacking the AP sites’ open-ring aldehyde form^24–26^. The resulting Schiff-base intermediate is then converted into a thiazolidine ring upon reaction with the sulfhydryl group of Cys2 (Figure 1B).

**Figure 1.**
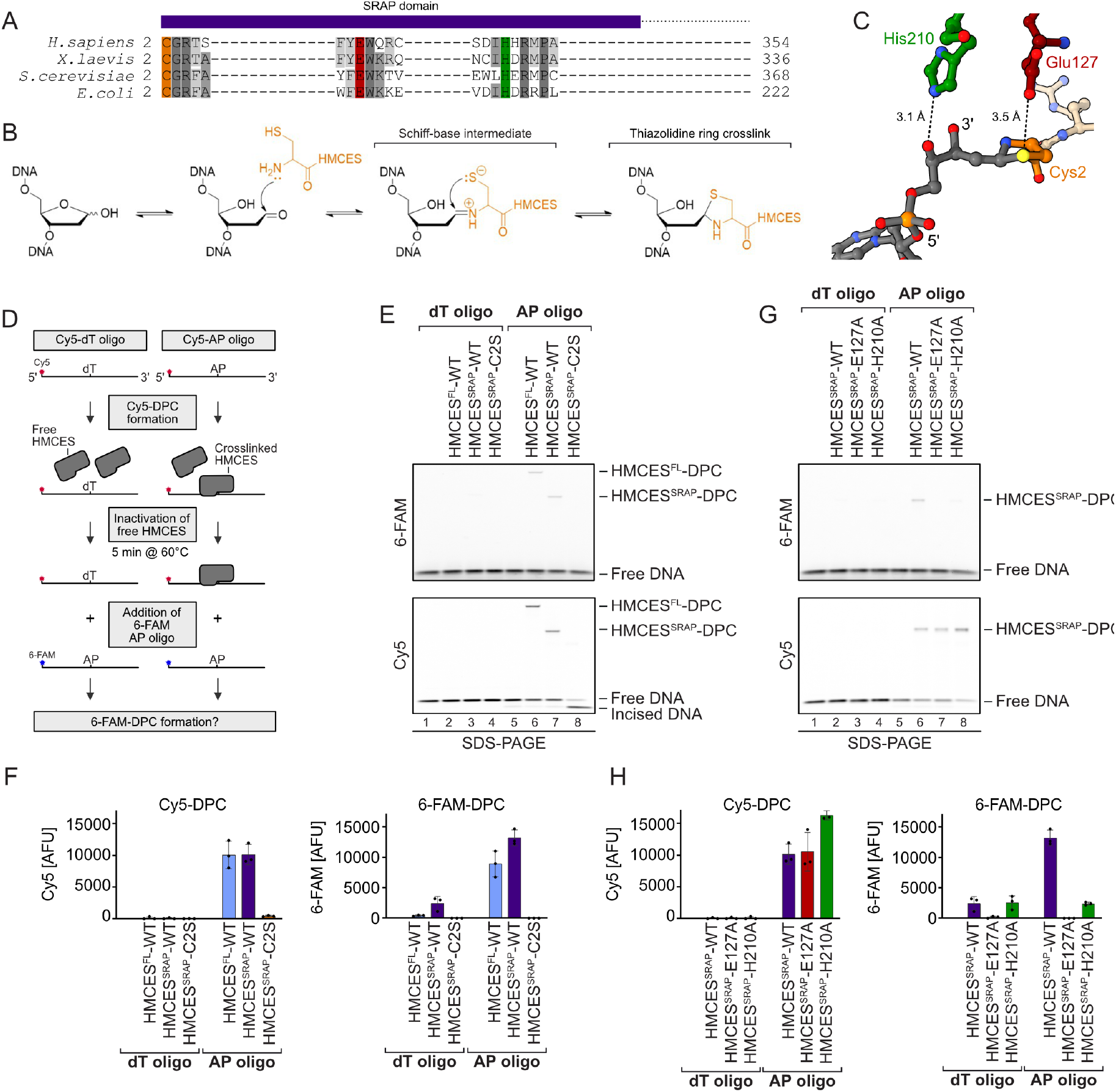
HMCES-DNA-protein crosslinks are reversible. **(A)** SRAP domain sequence alignment highlighting key active side residues in *H. sapiens, X. laevis., S. cerevisiae*, and *E. coli* HMCES homologs (Cys2 = orange, Glu127 = red, His210 = green). **(B)** Proposed reaction mechanism of SRAP domain crosslinking to an AP site. **(C)** Crystal structure of HMCES’ active site crosslinked to an AP site (PDB: 6OE7). DNA is shown in grey. Active site residues are coloured as in (A). Interatomic distances (Å) are labelled. **(D)** Schematic of the assay shown in (E) and (G). HMCES^FL^ and HMCES^SRAP^ (WT or active site variants) were incubated for 1 h at 37°C with a Cy5-labelled 30mer oligonucleotide containing either a dT or an AP site at position 15. Afterwards, non-crosslinked HMCES was inactivated by heat denaturation at 60°C for 5 min. A second 6-FAM-labelled 30mer oligonucleotide containing an AP site was added and formation of 6-FAM DPCs was assessed. **(E)** HMCES^FL^- and HMCES^SRAP^-WT and HMCES^SRAP^-C2S-DPC formation with Cy5- and 6-FAM-oligonucleotides were analysed using denaturing SDS-PAGE. **(F)** Quantification of DPC formation shown in (E), left panel: DPC formation to 6-FAM oligonucleotide, right panel: DPC formation to Cy5 oligonucleotide. Bar graph shows the mean of three independent experiments ± SD. **(G)** HMCES^SRAP^-WT and variants DPC (E127A or H210A) formation with Cy5- and 6-FAM-oligonucleotides were analysed using denaturing SDS-PAGE. **(H)** Quantification of DPC formation shown in (G), left panel: DPC formation to 6-FAM oligonucleotide, right panel: DPC formation to Cy5 oligonucleotide. Bar graph shows the mean of three independent experiments ± SD.

The protective function of HMCES-DPCs is particularly important, when cells face substantial amounts of AP sites, for example upon exposure to genotoxic agents^27^, overexpression of the cytosine deaminase APOBEC3A^23,28^, or AID-induced somatic hypermutation^29^. In addition, HMCES-DPCs were shown in *Xenopus* egg extracts to arise as intermediates of replication-coupled DNA inter-strand crosslink (ICL) repair^22^. Unhooking of the ICL by a DNA glycosylase yields an AP site, to which HMCES crosslinks. In egg extracts, HMCES-DPCs are mainly degraded by the SPRTN protease^22^, which requires unfolding of the protein adduct by the FANCJ helicase^30^. While human SPRTN cleaves HMCES-DPCs *in vitro* if FANCJ is present^30^, it is unclear to what extent SPRTN is required for repair in mammalian cells, where proteasomal HMCES-DPC degradation has been reported^17^. Notably, HMCES-DPCs are lost over time in egg extracts even if SPRTN is depleted and proteasomal activity inhibited^22^, indicating the existence of an additional mechanism that resolves HMCES-DPCs. Recent work indicated that SRAP-DPCs can undergo reversal in principle^31^, but it remained unclear how reversal can be reconciled with the need to protect AP sites in ssDNA.

Here, we use *in vitro* reconstitution and experiments in *Xenopus* egg extracts to dissect the principles of a non-proteolytic release mechanisms for HMCES-DPCs. We demonstrate that DPC release is determined by DNA context and occurs in two steps. First, a conserved glutamate residue located in HMCES’ active site catalyses the reversal of the thiazolidine crosslink. Second, HMCES either re-crosslinks, if affinity to the underlying DNA structure is high, or releases the AP site, if affinity is low. As a consequence, HMCES efficiently protects AP sites in ssDNA and at ssDNA-dsDNA junctions but releases them once the DPC is bypassed by the replication machinery and transferred into dsDNA.

## Results

### HMCES-DNA-protein crosslinks are reversible

Once HMCES-DPCs form, they appear stable over several days at room temperature *in vitro^24^*. To test whether HMCES remains irreversibly attached during incubation or constantly cycles between a crosslinked and a non-crosslinked state, we designed an assay to assess the reversibility of HMCES-DPCs (Figure 1D, schematic). First, we generated AP sites by incubating a Cy5-labelled 30mer DNA oligonucleotide containing a deoxyuridine (dU) at position 15 with uracil-DNA glycosylase (UDG). DNA containing dT instead of dU served as a control. DPCs were then generated by addition of recombinant full-length HMCES (HMCES^FL^) or the catalytic SRAP domain (HMCES^SRAP^). Next, reactions were exposed to a short heat treatment (5 min, 60°C), which inactivates free HMCES while not affecting crosslinked HMCES^30^. Finally, a 6-FAM-labelled AP site-containing DNA oligonucleotide was added to all reactions to test whether HMCES can be released from the Cy5-oligonucleotide and re-crosslink to the 6-FAM-oligonucleotide. Indeed, we observed formation of DPCs between 6-FAM-labelled DNA and HMCES^FL^ and HMCES^SRAP^ (Figure 1E, lanes 6 and 7, 1F, and S1A), suggesting that some DPCs between HMCES and the Cy5-oligonucleotide reverted which in turn allowed re-crosslinking to the 6-FAM-oligonucleotide. 6-FAM-DPCs did not form if a Cy5-dT-oligonucleotide was used (Figure 1E, lanes 2 and 3), indicating that inactivation of free HMCES was efficient, or if HMCES’ catalytic cysteine was replaced by serine (HMCES^SRAP^-C2S) (Figure 1E, lane 8).

Next, we asked whether HMCES-DPC reversal occurs spontaneously or whether it is an enzymatic process. The active site of HMCES features, in addition to the catalytic cysteine at position 2, two highly conserved amino acid residues, Glu127 and His210 (Figure 1A and 1C). Structural data suggest that both residues stabilize the transient Schiff-base intermediate during DPC formation^24–26^. Nonetheless, substitution of the corresponding glutamate residue in the prokaryotic HMCES ortholog YedK results in only reduced DPC formation^24,26^, while the effect of substituting the histidine remains controversial with reports of decreased and increased DPC formation^24,26^. Consistently, we observed that human HMCES^SRAP^ with substitution of Glu127 (E127A) or His210 (H210A) were able to form DPCs with Cy5-labelled AP site-containing DNA in our assay (Figure 1G, lanes 7 and 8, Cy5 scan, 1H, S1B, and S1C). However, HMCES^SRAP^-E127A and -H210A variants did not form DPCs with the subsequently added 6-FAM-oligonucleotide (Figure 1G, lanes 7 and 8, 6-FAM scan, and 1H). These results suggest that stabilisation of the Schiff-base intermediate by Glu127 and His210 is not essential for DPC formation *per se* but may rather be important to reverse thiazolidine ring formation, perhaps explaining the strict conservation of both residues during evolution. In agreement with a recent study^31^, we conclude that HMCES-DPCs are reversible, that released HMCES retains the ability to re-crosslink, and that release is an enzymatic process requiring conserved active site residues.

### Release of HMCES-DPCs is determined by DNA context

The fact that HMCES-DPCs are reversible raises the question whether the release is regulated. AP sites must be protected in ssDNA to prohibit strand breakage, but HMCES-DPC formation may be less favourable in dsDNA, where it would prohibit initiation of AP site repair by AP endonucleases. In line with the need to stabilize AP sites in ssDNA, DPC formation by HMCES^SRAP^-WT occurs efficiently in ssDNA and at ssDNA-dsDNA junctions with a 5’-flap (Figure 2A, 2B, and refs^17,24^). In contrast, DPC formation does not occur in dsDNA (Figure 2A, 2B and ref^17^), because HMCES cannot accommodate AP sites in its active site if dsDNA is present on the 3’-site of the lesion^24^. Substitution of Glu127 and His210 had no effect on the specificity of DPC formation, but HMCES^SRAP^-E127A crosslinked slower (Figure 2A and 2B), which may be related to a recently proposed role for Glu127 in AP site ring opening^31^. To understand whether DNA context also influences DPC release, we first generated DPCs between HMCES^SRAP^ and an AP site in ssDNA before annealing complementary reverse oligonucleotides to generate either a ssDNA-dsDNA junction or fully dsDNA (Figure 2C, schematic). Strikingly, HMCES^SRAP^-DPCs were stable in ssDNA or ssDNA-dsDNA junctions but reversed quickly in dsDNA (Figure 2C and 2D). In contrast, HMCES^SRAP^-variants H210A and E127A were partially (H210A) or entirely (E127A) defective for reversal in dsDNA (Figure 2C and 2D). HMCES^SRAP^-E127A was previously reported to display increased DNA binding^26^, which we confirmed using fluorescence polarization (Figure S2A) and electrophoretic mobility shift assays (Figure S2B). To test whether the inability of this variant to reverse is related to increased DNA binding, we combined E127A with a substitution of Arg98 (R98E), which is located within the HMCES^SRAP^-ssDNA interface^25^. DPC formation and release was not affected by the R98E substitution (Figure 2A-D, S1B and S1C), despite severely reduced DNA binding activity (Figure S2A, S2B, and ref^17^). In combination with E127A, substitution of Arg98 decreased DNA binding below WT levels (Figure S2A and S2B) but did not restore the ability to revert the crosslink (Figure S2C and S2D). Thus, the reversal defect of HMCES^SRAP^-E127A-DPCs is unrelated to increased DNA binding affinity. We conclude that DPC release is not only an active process requiring Glu127 (and partially His210) but is also determined by DNA context. DPC release displays opposite specificity to DPC formation, which correlates with the biological need to protect AP sites in ssDNA but not in dsDNA.

**Figure 2.**
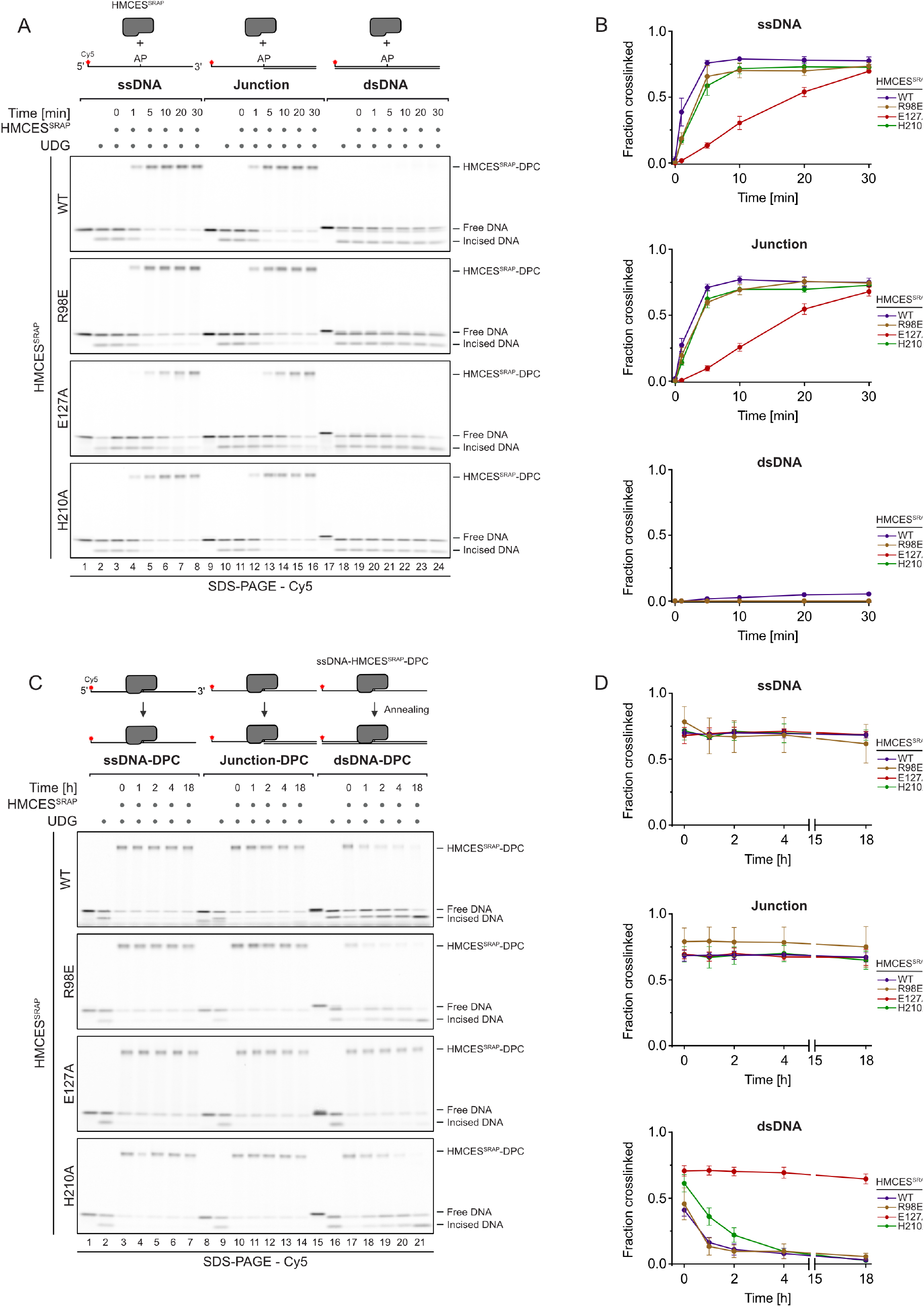
Release of HMCES-DPCs is determined by DNA context. **(A)** Kinetics of DPC formation by HMCES^SRAP^ (WT, R98E, E127A, or H210A variants) to ssDNA, junction DNA, and dsDNA. Corresponding reverse oligonucleotides were annealed to ssDNA to create DNA junction and dsDNA prior to adding HMCES^SRAP^. To ssDNA, a non-complementary oligonucleotide was added as control. HMCES^SRAP^-WT and variants were incubated with different DNA structures for the indicated amount of time at 37°C prior to separation by denaturing SDS-PAGE. **(B)** Quantification of DPC formation assays shown in (A): data represent the mean of three individual experiments ± SD. **(C)** DPC reversal kinetics of indicated variants in ssDNA, DNA junction, and dsDNA. DPCs were pre-formed in ssDNA before corresponding reverse oligonucleotides were annealed (for ssDNA reactions a non-complementary oligonucleotide was added). DPC reversal was then monitored after incubation for the indicated amount of time at 37°C using denaturing SDS-PAGE. **(D)** Quantification of DPC reversal of HMCES^SRAP^-WT and variants shown in (C): data represent the mean of three individual experiments ± SD. See also Figure S2.

### Release of HMCES-DPCs is determined by binding affinity to the underlying DNA

Next, we wanted to understand how DNA context controls the release of HMCES^SRAP^-DPCs. Our results so far could be explained by a model in which all HMCES-DPCs constantly revert independent of DNA context and that specificity is only determined by HMCES’ ability to reform the crosslink after release, which does not occur in dsDNA. However, it remained unclear how HMCES could efficiently protect AP sites, if it would constantly dissociate from the lesion. To gain more detailed insights into DPC reversal in different DNA structures, we first generated HMCES^SRAP^-DPCs in ssDNA and at ssDNA-dsDNA junctions (DPCs in dsDNA released too quickly to be assessed by this assay). We then added HMCES^FL^ in ten-fold excess to outcompete HMCES^SRAP^ upon release of the AP site (Figure 3A and 3B, schematic). This set-up allowed us to evaluate HMCES^SRAP^-DPC reversal by monitoring the appearance of HMCES^FL^-DPCs over time. Notably, HMCES^SRAP^-DPCs were released over time in ssDNA (Figure 3A, lanes 3-8) but were much more stable at ssDNA-dsDNA junctions (Figure 3B, lanes 3-8); DPCs formed by the E127A variant were not released in either setting (Figure 3A, lanes 15-20 and Figure 3B, lanes 15-20). We wondered whether the enhanced release of WT DPCs from ssDNA was related to the previously reported preferential binding of the SRAP domain to ssDNA-dsDNA junctions compared to ssDNA^24^. Accordingly, we hypothesized that the active site of HMCES may constantly cycle between a crosslinked and a non-crosslinked state independent of DNA context, but that actual dissociation from the underlying DNA substrate would in addition be determined by binding affinity. To test this idea, we asked whether the reduced DNA binding affinity of HMCES^SRAP^-R98E would affect reversal. Indeed, HMCES^SRAP^-R98E-DPCs reversed much more rapidly in ssDNA and at ssDNA-dsDNA junctions than WT-DPCs (Figure 3A, lanes 9-14 and Figure 3B, lanes 9-14). Taken together, these results suggest that HMCES-DPC release is governed by two major components. First, the principal capacity of Glu127 to catalyse reversal ensures cycling of the active site between a crosslinked and a non-crosslinked state. Second, the binding strength to the underlying DNA structure then determines whether HMCES re-crosslinks or dissociates while in the non-crosslinked state, which occurs if affinity is low (e.g., within dsDNA or in context of R98E-DPCs).

**Figure 3.**
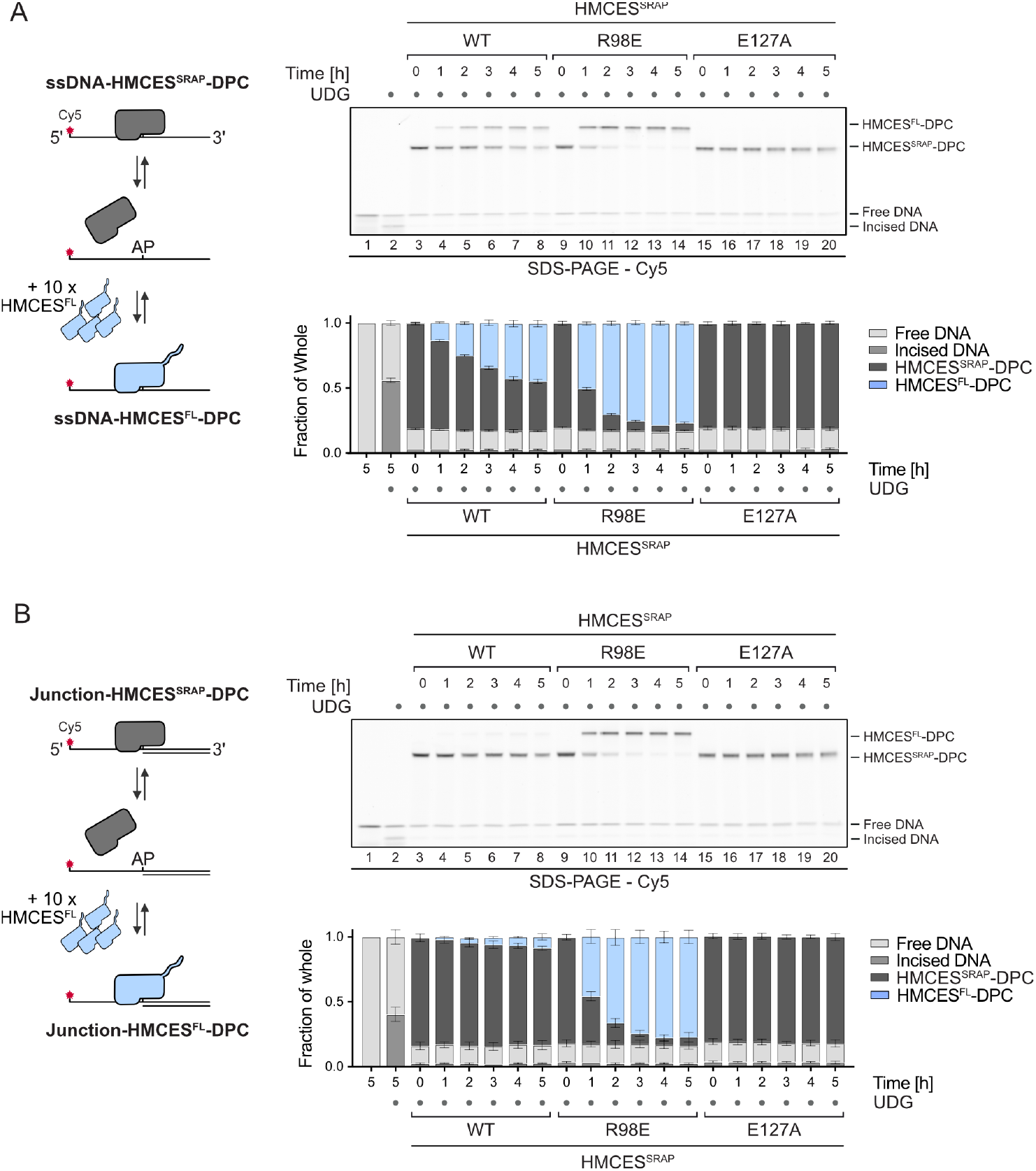
Release of HMCES-DPCs is determined by binding affinity to the underlying DNA. **(A-B)** Competition assay between HMCES^FL^ and indicated HMCES^SRAP^ variants. HMCES^SRAP^-DPCs in ssDNA (A) or at ssDNA-dsDNA junctions (B) were pre-formed and then incubated with ten-fold excess of HMCES^FL^ for the indicated amount of time at 37°C prior to separation by denaturing SDS gel (upper panels). Quantification of competition assay: bar graphs show the mean of three independent experiments ± SD (lower panels).

### Release of HMCES-DPCs restricts crosslink formation to physiologically relevant situations

Next, we wanted to understand how DPC release relates to HMCES’ ability to block APE1 endonuclease from incising AP sites. APE1 efficiently cleaves AP sites at ssDNA-dsDNA junctions and in dsDNA, but shows little activity in ssDNA (Figure S3A and ref^32^). Therefore, we generated HMCES^SRAP-^WT, -R98E and -E127A-DPCs at ssDNA-dsDNA junctions and in dsDNA and incubated them with APE1. As reported previously^17^, WT-DPCs shielded AP sites from APE1 incision at ssDNA-dsDNA junctions (Figure 4A, lanes 8-10). In contrast, R98E-DPCs failed to protect against APE1 (Figure 4A, lanes 14-16), suggesting that the increased release of this variant compromises its ability to protect AP sites against APE1 incision. In dsDNA, both WT- and R98E-DPCs did not prevent AP site cleavage (Figure 4B, lanes 8-10 and 14-16, respectively), while E127A-DPCs fully blocked incision (Figure 4B, lanes 20-22). Thus, indicating that failure to auto-release results in inhibition of AP site repair in dsDNA. Collectively, these data show that HMCES-DPC release must be finely balanced to (1) ensure protection of AP sites at ssDNA-dsDNA junctions against potentially catastrophic APE1 incisions (which is compromised upon hyper-reversal in the HMCES^SRAP^-R98E variant) and to (2) allow deprotection of AP site in dsDNA so that APE1 can initiate repair (which is compromised upon hypo-reversal in the HMCES^SRAP^-E127A variant).

**Figure 4.**
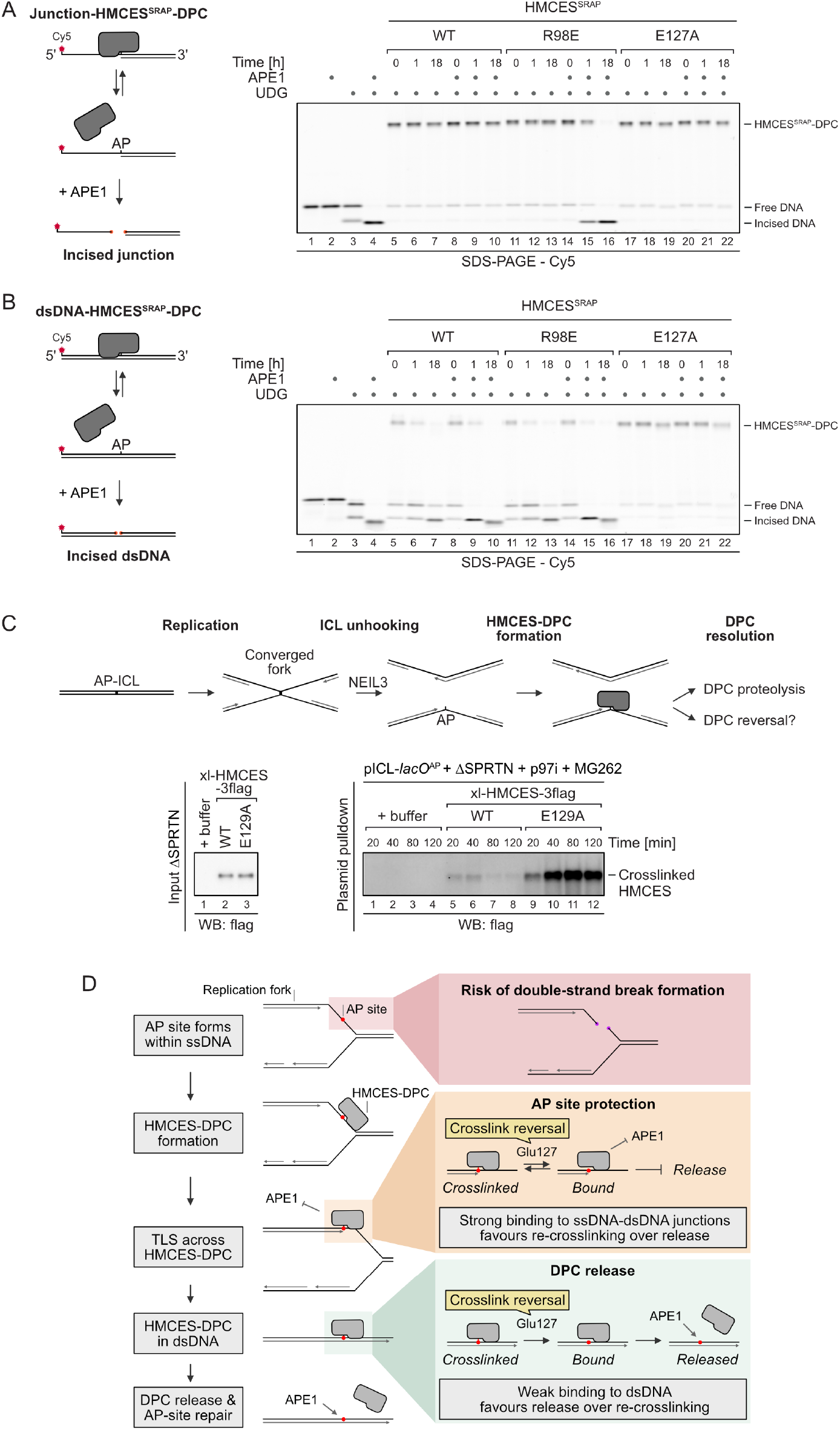
Auto-release of HMCES-DPCs restricts crosslink formation to physiologically relevant situations. **(A-B)** APE1 incision of an AP site protected by the indicated HMCES^SRAP^-DPC variants at ssDNA-dsDNA junctions (A) or within dsDNA (B). Free dU-containing DNA was incubated alone or in the presence of UDG and HMCES^SRAP^ for 1 h at 37°C. Next, corresponding reverse oligonucleotides were annealed to generate a ssDNA-dsDNA junction (A) or dsDNA (B), and reactions were incubated alone or with APE1 for the indicated amount of time at 37°C prior to separation by denaturing SDS-PAGE. **(C)** Upper panel: depiction of AP-ICL repair by NEIL3 in *Xenopus* egg extract. ICL unhooking by NEIL3 generates an AP site that in turn reacts with HMCES to form a DPC. The resulting HMCES-DPC can either be removed by proteolysis or by catalyzed crosslink reversal. Lower left panel: the extracts used in the replication reactions shown in the lower right panel were blotted for 3flag epitope. Lower right panel: pICL-*lacO*A^P^ was replicated in the indicated egg extracts supplemented with p97i, MG262, and recombinant xl-HMCES-3flag. Chromatin was recovered under stringent conditions, the DNA was digested, and released proteins were separated by SDS-PAGE. Xl-HMCES-3flag was detected by blotting for 3flag epitope. **(D)** Model of AP site protection by coordinated formation and release of HMCES-DPCs. See also Figure S3.

### Release of HMCES-DPCs during replication-coupled ICL repair

Next, we wanted to test whether auto-release contributes to the resolution of physiological HMCES-DPCs that occur as intermediates during replication-coupled ICL repair^22^. In *Xenopus* egg extracts, we monitored replication of plasmids containing an AP-ICL, a lesion that forms when an AP site reacts with a nucleobase of the opposing DNA strand forming a covalent crosslink^33^. Such crosslinks are targeted by two replication-coupled repair mechanisms^34^. They can be repaired by the Fanconi anemia pathway, which we prevented by blocking the unloading of CMG helicase by chemical inhibition of p97 (p97i)^35,36^. Alternatively, AP-ICLs can be unhooked by the NEIL3 glycosylase^35^, which yields an AP site leading to formation of a HMCES-DPC (Figure 4C, schematic, and ref^22^). To investigate auto-release of these DPCs, we examined repair of an AP-ICL containing plasmid (pICL-*lacO*A^P^) using egg extracts supplemented with WT or E129A recombinant C-terminally 3flag-tagged *Xenopus laevis* HMCES (xl-HMCES-3flag) (Figure 4C, left panel); the *Xenopus* HMCES E129A variant corresponds to the human E127A variant and was observed to both slow DPC formation at a ssDNA AP site and fully block DPC release from dsDNA, as observed for human proteins (Figure S3B and S3C). In addition, we prevented DPC proteolysis in the extract by immunodepletion of SPRTN and inhibition of the proteasome using MG262 (Figure S3D). Using this set-up, we then monitored replication of the AP-ICL-containing plasmid, which was comparable in all conditions tested (Figure S3E). In parallel, we monitored DPC formation using stringent plasmid pulldown followed by DNA digestion and western blotting. As described previously for endogenous HMCES^22^, we observed formation of tagged HMCES-WT-DPCs after 20 minutes (correlating with ICL unhooking by NEIL3; Figure S3C) and these DPCs mostly resolved by 80 minutes (Figure 4C, lanes 5-8). In contrast, HMCES-E129A-DPCs were not resolved and accumulated to dramatic levels that persisted to 120 minutes (Figure 4C, lanes 9-12). These data strongly suggest that DPC reversal catalysed by Glu129 is solely responsible for the release of HMCES-DPCs during replication-coupled ICL repair if proteolysis is inhibited.

## Discussion

In this study, we found that HMCES-DPCs do not necessarily require proteolytic repair because they feature a built-in release mechanism. Our data suggest a model in which auto-release of HMCES-DPCs occurs in two distinct steps (Figure 4D). First, the conserved Glu127 residue (with a minor contribution of His210) catalyses the reversal of the crosslink between HMCES’ active site cysteine and the AP site, as also observed in other recent work^31^. Second, the cysteine then either re-crosslinks or HMCES dissociates from DNA resulting in release of the AP site. The decision between these two options appears to be determined by binding strength to the underlying DNA. HMCES binds tightly to ssDNA-dsDNA junctions, which favours re-crosslinking over release. In contrast, HMCES binds poorly to dsDNA, resulting in release. Thus, this model explains how HMCES can protect AP sites at ssDNA-dsDNA junctions against incisions by AP endonucleases, while promoting AP site cleavage within dsDNA.

HMCES-DPCs form specifically in ssDNA contexts^17^, which leads to the question as to how HMCES-DPCs are transferred to dsDNA, where release could occur. Two settings seem plausible; one, TLS polymerases extend nascent strands across intact HMCES-DPCs, as has been observed in a FANCJ-dependent manner in *Xenopus* egg extracts and *in vitro^30^;* two, nascent strands may be extended upon template switching, which would avoid the need for TLS and thus ensure error-free repair^17,23,27^. Notably, in either case, extension of the nascent strand would prevent DPC proteolysis by SPRTN, which requires the presence of a ssDNA-dsDNA junction in close proximity to the protein adduct to become activated^8,37^.

The relative importance of release, which has the added benefit of recycling the enzyme, versus proteolytic repair of HMCES-DPCs is an interesting future question. HMCES^E127A^ has been reported to complement the sensitivity of *HMCES* KO cells to AP site-inducing drugs^27^ and to protect AP sites during somatic hypermutation^29^, which is in line with this variant’s ability to form DPCs. We have not observed toxicity in human cells upon over-expression of HMCES^E127A^ (data not shown), which may indicate that proteolytic repair fully compensates for the lack of DPC release. Both proteases targeting HMCES-DPCs, SPRTN and the proteasome, are essential for cell viability, which prohibits the analysis of HMCES^E127A^ toxicity in the absence of DPC proteolysis in cells. Thus, the effects of defective auto-release remain unclear. However, human cells expressing HMCES^R98E^ are sensitive to ionising radiation^17^, which together with our data shows that increased DPC release compromises HMCES ability to protect cells against AP-sites in ssDNA. The precise regulation of the HMCES-DPC auto-release mechanism identified in this study emphasizes the need to restrict DPC formation to an absolute minimum.

## Acknowledgements

We thank T. Fröhlich for mass spectrometry analyses of recombinant HMCES protein and T. Mackens-Kiani for help with preparing Figure 1C. S.D. is supported by the International Max-Planck Research School for Molecular Life Sciences. Research in the lab of D.R.S is supported by NIH grant no. GM129422. Research in the lab of J.S. is supported by the European Research Council under the European Union’s Horizon 2020 research and innovation program (grant agreement number 801750), by the Alfried Krupp Prize for Young University Teachers awarded by the Alfried Krupp von Bohlen und Halbach Foundation, the European Molecular Biology Organization (YIP4644), the Vallee Foundation, and by the Deutsche Forschungsgemeinschaft (DFG, German Research Foundation) (Project ID 213249687 – SFB 1064).

## Author Contributions

M.D. and J.S. conceived the project. M.D. and S.D. performed the majority of experiments. K.T.N performed experiments in Xenopus egg extracts supervised by D.R.S.. F.G. performed experiments shown in Figure S1B and S1C. D.Y. performed replicates of several experiments. M.D., S.D., and J.S. wrote the manuscript with input from all authors. J.S. acquired funding and supervised the work.

---

**Figure S1.**
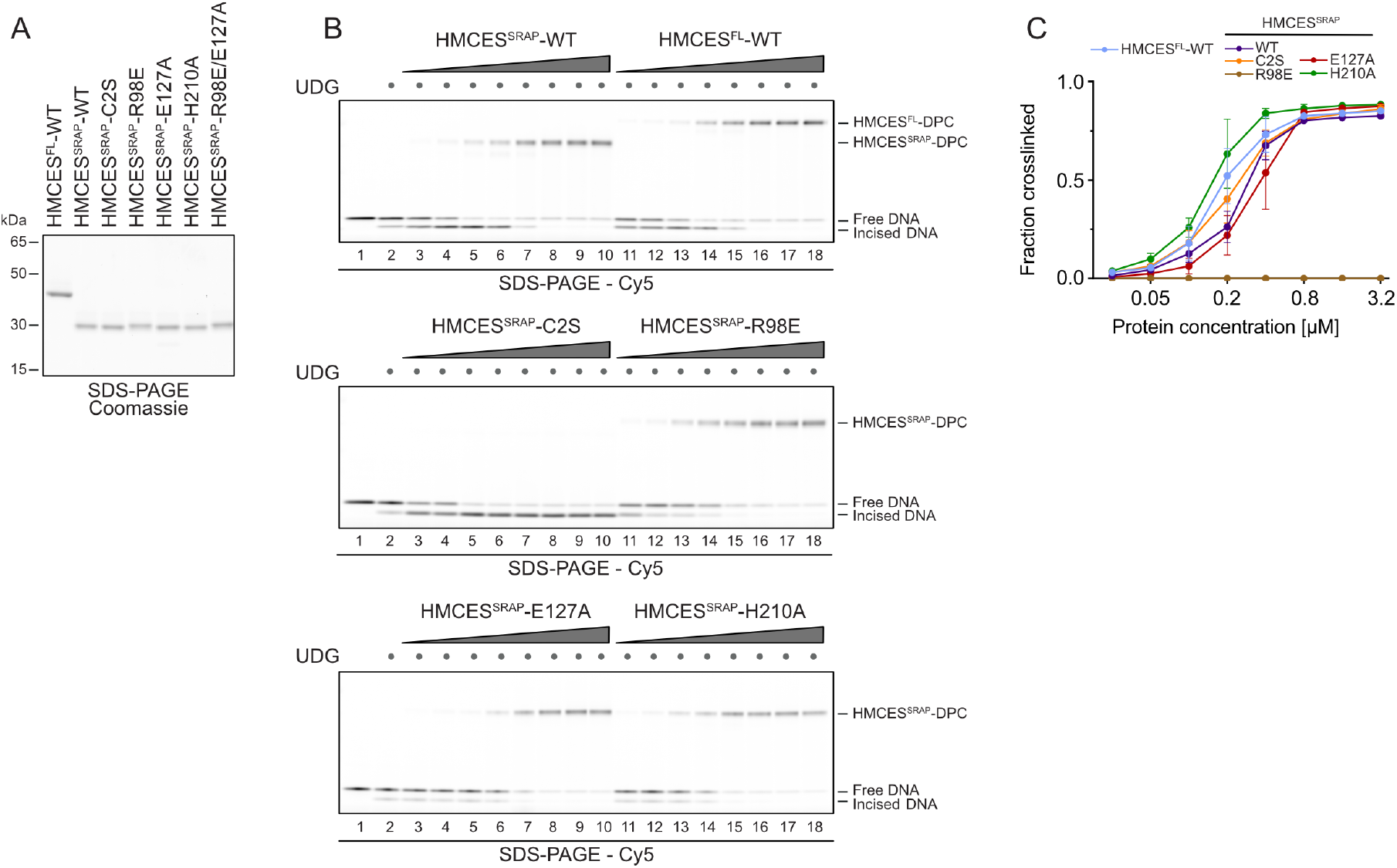
HMCES-DNA-protein crosslinks are reversible. Related to Figure 1 **(A)** Coomassie stained SDS-PAGE gel showing recombinant purified human HMCES^FL^-WT, HMCES^SRAP^-WT, HMCES^SRAP^-C2S, HMCES^SRAP^-R98E, HMCES^SRAP^-E127A, HMCES^srap^-H210A, HMCES^SRAP^-R98E/E127A proteins used in this study. **(B)** DPC formation of HMCES^FL^, HMCES^SRAP^ (WT or indicated variants). dU-containing DNA (0.1 μM) was incubated alone or with UDG and increasing concentrations of HMCES (0.025 μM, 0.05 μM, 0.1 μM, 0.2 μM, 0.4 μM, 0.8 μM, 1.6 μM, 3.2 μM), as indicated for 1 h at 37°C prior to analysis by denaturing SDS-PAGE. **(C)** Quantification of HMCES-DPC formations shown in (B): data represent the mean of three individual experiments ± SD.

**Figure S2.**
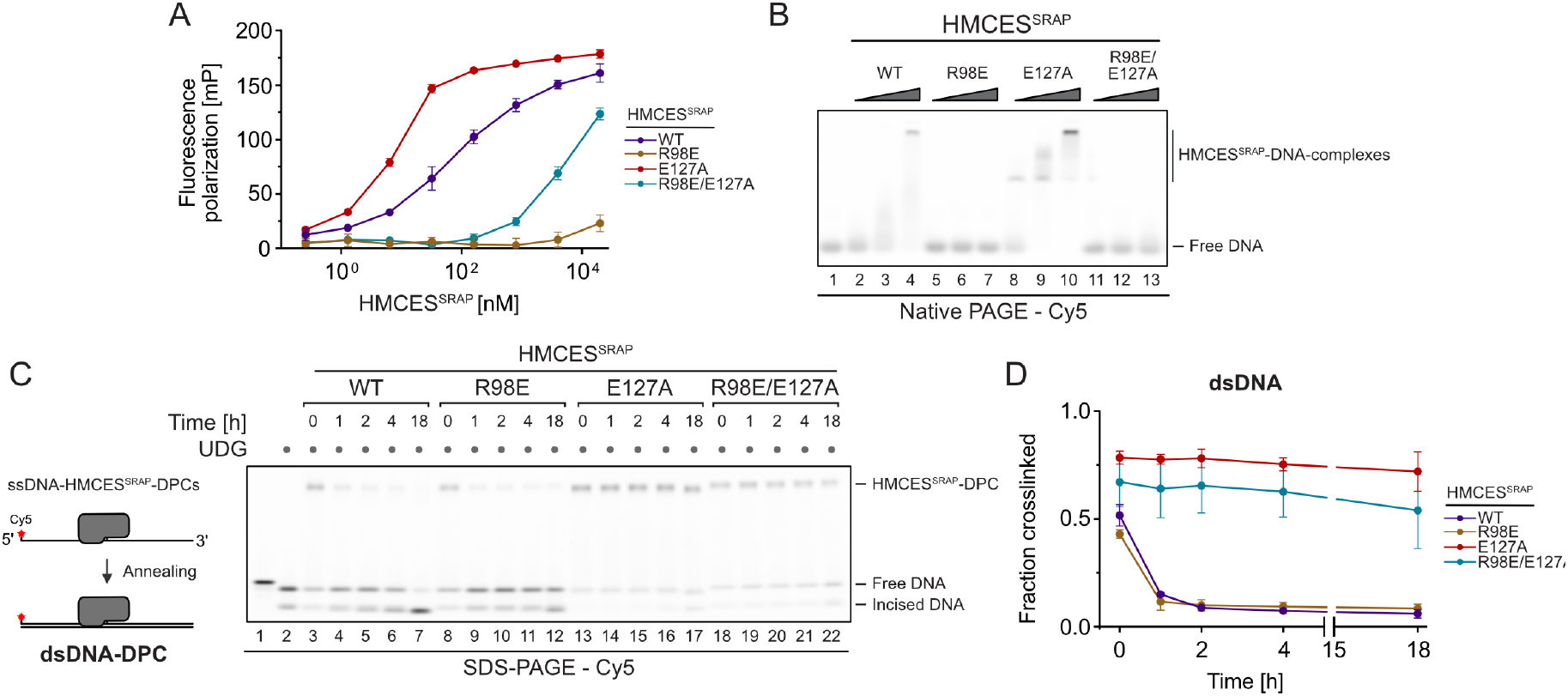
Release of HMCES-DPCs is determined by DNA context. Related to Figure 2 **(A)** Fluorescence polarization measurements of Cy5-labelled ssDNA (25 nM) incubated with increasing concentrations of HMCES^SRAP^-WT, HMCES^SRAP^-R98E, HMCES^SRAP^-E127A, or HMCES^SRAP^-R98E/E127A for 20 min on ice prior to measuring fluorescence polarization. **(B)** Non-covalent DNA binding of indicated HMCES^SRAP^ variants was assessed by electrophoretic mobility shift assay. A Cy5-labelled 30-mer ssDNA (0.1 μM) was incubated with HMCES^SRAP^ (0, 0.125, 0.5, or 2 μM) for 20 min at 4°C prior to analysis by native PAGE. **(C)** DPC reversal kinetics of indicated HMCES^SRAP^ variants in dsDNA. A corresponding reverse oligonucleotide was annealed to HMCES^SRAP^-DPCs, before incubation for the indicated amount of time at 37°C prior to separation by denaturing SDS-PAGE. **(D)** Quantification of DPC reversal kinetics shown in (C): data represent mean of three independent experiments ± SD.

**Figure S3.**
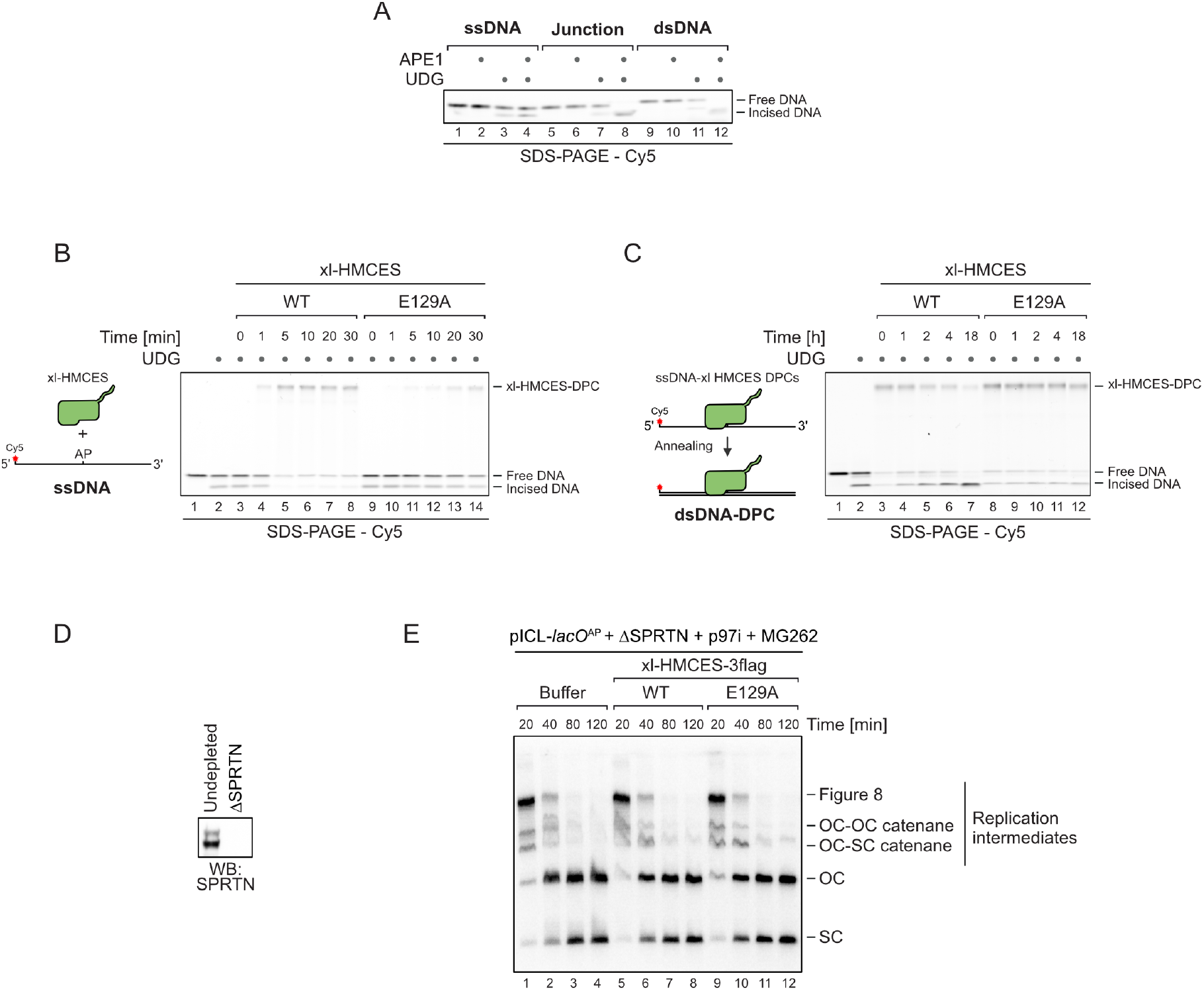
Auto-release of HMCES-DPCs restricts crosslink formation to physiologically relevant situations. Related to Figure 4 **(A)** APE1 incision of an AP site in ssDNA, DNA junction, and dsDNA. A Cy5-labelled 30-mer ssDNA was incubated alone or with UDG for 1 h at 37°C. Corresponding reverse oligonucleotides for DNA junction or dsDNA were annealed (For ssDNA a non-complementary oligonucleotide was added). Next, samples were incubated with APE1 for 18 h at 37°C before separation by denaturing SDS-PAGE. **(B)** Kinetics of DPC formation by xl-HMCES-WT and -E129A to ssDNA. A non-complementary oligonucleotide was added to match the experimental set-up to that in Figure 2A. xl-HMCES-WT and -E129A were incubated with ssDNA for the indicated amount of time at 37°C prior to separation by denaturing SDS-PAGE. **(C)** Kinetics of DPC reversal kinetics of xl-HMCES-WT and -E129A in dsDNA. DPCs were pre-formed in ssDNA before a corresponding reverse oligonucleotide was annealed. DPC reversal was then monitored after incubation for the indicated amount of time at 37°C using denaturing SDS-PAGE. **(D)** The extract used in the replication reactions shown in Figure 4C was immunoblotted for SPRTN. **(E)** In parallel with the reactions shown in Figure 4C, pICL-*lacO*^AP^ was replicated in the indicated egg extracts supplemented with p97i, MG262, recombinant xl-HMCES-3flag, and [a-^32^P]dCTP. Replication intermediates were separated on a native agarose gel and visualized by autoradiography. SC, supercoiled. OC, open circular.

## Methods

### RESOURCE AVAILABILITY

#### Lead Contact

Further information and requests for resources and reagents should be directed to and will be fulfilled by the Lead Contact, Julian Stingele.

#### Materials Availability

All plasmids are available on request.

#### Data and Code Availability

- Original gel images reported in this paper will be shared by the lead contact upon request.
- This study did not generate original code.
- Any additional information required to reanalyse the data reported in this paper is available from the lead contact upon request.

### EXPERIMENTAL MODEL AND SUBJECT DETAILS

#### Xenopus laevis

*Xenopus laevis* female frogs (Xenopus1, Cat# 4280; age > 2 years) were used for collecting eggs. To induce ovulation, female frogs were injected with human chorionic gonadotropin (hCG). Males (Xenopus1, Cat# 4290, age > 1 year) were used for collecting sperm chromatin. All frogs were maintained in the pathogen-free amphibian facility in the CCE Department at Caltech in compliance with IACUC approval (Protocol IA20-1797, approved 28 May 2020). The Caltech has an approved Animal Welfare Assurance (no. D16-00266) from the NIH Office of Laboratory Animal Welfare.

### METHOD DETAILS

#### Protein expression and purification

##### HMCES^SRAP^

An open reading frame containing human HMCES^SRAP^ (amino acids 1-270) was codon-optimized for bacterial expression and cloned in frame with a C-terminal His6-tag in a pNIC plasmid. HMCES^SRAP^-variants (-C2S, -R98E, -E127A, -H210A, -R98E/E127A) were generated by introducing point mutations using the Q5 site-directed mutagenesis kit (New England BioLabs), following manufacturer’s instructions. Mutations were confirmed by Sanger sequencing. BL21 (DE3) *Escherichia coli* cells were transformed with the corresponding plasmids for protein expression. Cells were grown in Terrific broth (TB) medium at 37°C to OD_600_ 0.7. Protein expression was induced by addition of 0.5 mM isopropyl-β-D-thiogalactoside (IPTG) for 4 h. Cells were harvested, snap-frozen in liquid nitrogen and pellets were stored at −80°C. For protein purification, cells were resuspended in buffer A (50 mM HEPES/KOH pH 7.8, 500 mM KCl, 5 mM MgCl_2_, 30 mM imidazole, 10% glycerol, 0.1% IGEPAL, 0.04 mg/mL Pefabloc SC, cOmplete EDTA-free protease inhibitor cocktail tablets, 1 mM tris(2-carboxyethyl)phosphine hydrochloride (TCEP)) and lysed by sonication. All subsequent steps were carried out at 4°C. Cell lysates were incubated with Benzonase nuclease (45 U/mL lysate) for 30 min, before cell debris was removed by centrifugation at 18,000 g for 30 min. Cleared and filtered supernatants were applied to 3 mL Ni-NTA Agarose (QIAGEN) equilibrated in buffer B (20 mM HEPES/KOH pH 7.8, 500 mM KCl, 5 mM MgCl_2_, 30 mM imidazole, 10% glycerol, 1 mM TCEP). Next, beads were washed with 15 column volumes (CV) of buffer B before protein was eluted in 2 CV of buffer C (20 mM HEPES/KOH pH 7.8, 500 mM KCl, 5 mM MgCl_2_, 300 mM imidazole, 10% glycerol, 1 mM TCEP). The elution was concentrated to 2 mL using a 10 kDa molecular weight cut-off Amicon Ultra centrifugal filter prior to loading on a HiLoad^®^ 16/600 Superdex^®^ 200 pg column equilibrated in buffer D (20 mM HEPES/KOH pH 7.8, 150 mM KCl, 5 mM MgCl_2_, 10% glycerol, 1 mM TCEP) for size exclusion chromatography. Eluted protein fractions were collected and concentrated with a 10 kDa molecular weight cut-off Amicon Ultra centrifugal filter. Concentrated protein was aliquoted, snap-frozen and stored at −80°C. Removal of the N-terminal methionine was confirmed by mass spectrometry.

##### HMCES^FL^

For full length HMCES (HMCES^FL^) the open reading frame was codon optimized and cloned in a pNIC plasmid in frame with a C-terminal TwinStrep-ZB-tag. Recombinant HMCES protein was expressed and purified using a protocol for purification of SPRTN ^37^, with small modifications to some buffers. Cell pellets were resuspended in buffer A (20 mM HEPES/KOH pH 7.5, 500 mM KCl, 5 mM MgCl_2_, 30 mM imidazole, 10% glycerol, 0.1% IGEPAL, 0.04 mg/mL Pefabloc SC, cOmplete EDTA-free protease inhibitor cocktail tablets, 1 mM TCEP). For washing steps buffer B (20 mM HEPES/KOH pH 7.5, 500 mM KCl, 5 mM MgCl_2_, 1 mM TCEP) was used. Protein was eluted from Strep-Tactin^®^XT Superflow^®^ high capacity cartridges with buffer B containing 50 mM Biotin and from HiTrap Heparin HP affinity columns in buffer C (20 mM HEPES/KOH pH 7.5, 1 M KCl, 5 mM MgCl_2_, 1 mM TCEP). For size exclusion chromatography and storage of the protein buffer D (20 mM HEPES/KOH pH 7.8, 150 mM KCl, 5 mM MgCl_2_, 10% glycerol, 1 mM TCEP) was used. Removal of the N-terminal methionine was confirmed by mass spectrometry.

##### Biotiny/ated LacI

Biotinylated LacI was purified as described previously^38^. Briefly, pET11a[LacR-Avi] and pBirAcm (Avidity) vectors were transformed into T7 Express competent cells. LacI and biotin ligase expression was induced with 1 mM IPTG in Luria-Bertani (LB) medium supplemented with 50 μM biotin for 2 h at 37°C. Cells were harvested, snap frozen and stored at −80°C. Cell pellets were lysed in lysis buffer (50 mM Tris-HCl pH 7.5, 5 mM EDTA, 100 mM NaCl, 1 mM DTT, 10% sucrose, cOmplete protease inhibitors, 0.2 mg/mL lysozyme, 0.1% Brij 58) for 30 min at room temperature (RT). Lysates were centrifuged at 21,300 g for 1 h at 4°C. Pellets containing chromatin-bound LacI was then suspended in 50 mM Tris-HCl pH 7.5, 5 mM EDTA, 1 M NaCl, 30 mM IPTG, 1 mM DTT and LacI was released from DNA by sonication followed by addition of polymin P to 0.03-0.06 % (w/v) at 4°C. Biotinylated LacI was precipitated with 37% ammonium sulfate, pelleted by centrifugation, and then suspended in 50 mM Tris-HCl pH 7.5, 1 mM EDTA, 2.6 M NaCl, 1 mM DTT and cOmplete protease inhibitors. Biotinylated LacI was then bound to SoftLINK avidin, washed with 50 mM Tris-HCl pH 7.5, 1 mM EDTA, 2.6 M NaCl, 1 mM DTT, cOmplete protease inhibitors, and eluted with 50 mM Tris-HCl pH 7.5, 100 mM NaCl, 1 mM EDTA, 5 mM biotin and 1 mM DTT. Pooled fractions containing biotinylated LacI were buffer exchanged into 50 mM Tris-HCl pH 7.5, 150 mM NaCl, 1 mM EDTA, 1 mM DTT using an Amicon Ultra-0.5 mL 3 kDa molecular weight cut-off filter unit. Biotinylated LacI was aliquoted, snap frozen and stored at −80°C.

##### USP2-cc

To purify USP2-catalytic core (USP2-cc), pH_10_E USP-cc plasmid was transformed into BL21 (DE3) *E. coli* cells. Expression was induced with 0.5 mM IPTG in LB medium for 16 h at 18°C. Cells were pelleted and lysed in lysis buffer E (20 mM Tris pH 8.0, 300 mM NaCl, 10 mM imidazole, 10% glycerol, 1% Triton X-100, 1 mg/mL lysozyme, cOmplete EDTA-free protease inhibitor cocktail tablets, 5 mM 2-β-mercaptoethanol (BME), 10 U/mL benzonase (Sigma, 70746-3). Cell lysates were cleared by centrifugation at 18,000 g for 20 min. His-tagged Usp2-cc was bound to Ni-NTA Agarose (QIAGEN) equilibrated in buffer F (20 mM Tris pH 8.0, 300 mM NaCl, 10 mM imidazole, 10% glycerol, 0.05% Triton X-100, 5 mM BME) for 4 h at 4°C, washed three times with buffer F, and then eluted with buffer G (20 mM Tris pH 8.0, 300 mM NaCl, 300 mM imidazole, 10% glycerol, 0.05% Triton X-100, 1 mM TCEP). Eluted protein was dialyzed in dialysis buffer H (20 mM Tris pH 8.0, 150 mM NaCl, 10% glycerol, 1 mM TCEP). Protein was then aliquoted, snap frozen and stored at −80°C.

##### xl-HMCES

An open reading frame containing *Xenopus laevis* HMCES (amino acid 2-336) was cloned in frame with a N-terminal His10-Ubiquitin (Ub) to generate pHUE-*xl*.HMCES plasmid^39^. pHUE-*xl*.HMCES(E129A) plasmid was generated by inverse PCR using primer pairs (5’-CAG GAC GGT GAA AAA CAA CCG TAC-3’/5’-GCG TTT CCA TGC ATA GAA CCC GTC C-3’). All constructs were confirmed by Sanger sequencing and transformed into ArcticExpress (DE3) competent cells for protein expression. Cells were grown in LB medium at 37°C to OD_600_ 0.6. Protein expression was induced by addition of 0.5 mM IPTG for 24 h. Cells were harvested, washed once with PBS (137 mM NaCl, 2.7 mM KCl, 10 mM Na_2_HPO_4_, 1.8 mM KH_2_PO_4_), snap-frozen and stored at −80°C. For protein purification, cells were resuspended in buffer E, incubated on ice for 30 min, and briefly sonicated. Cell debris was removed by centrifugation at 18,000 g for 20 min. Cleared supernatants were applied to Ni-NTA Agarose (QIAGEN) equilibrated in buffer F. Next, beads were washed thrice with 15 CV of buffer F and the protein was eluted with 4 CV of buffer G. The eluted protein was dialyzed against buffer H. The His-tagged proteins were incubated overnight at 4°C with His10-USP2-cc (molar ratio 1/100) to cleave the His10-Ub tag from the N-terminus of HMCES. The cleavage reaction mixtures were incubated with 1 mL pre-washed Ni-NTA agarose to remove His10-Ub, His10-USP2-cc and uncleaved His10-Ub-HMCES. HMCES in the flowthrough was further pufiried by anion exchange chromatograpy using mono Q50/5 GL column (Cytiva). Samples were eluted over a gradient of 150 to 100 mM NaCl. Fractions containing proteins were pooled and concentrated using Amicon Ultra-15 centrigufugal filter unit with 10 kDa molecular weight cut-off. Protein was aliquoted, snap-frozen and stored at −80°C.

##### xl-HMCES-3flag

To purify xl-HMCES-3flag and xl-HMCES(E129A)-3flag, a DNA sequence encoding 3flag was inserted downstream of xl-HMCES and xl-HMCES(E129A) in pHUE backbone plasmid to generate pHUE-xl.HMCES-3flag and pHUE-xl.HMCES(E129A)-3flag, respectively. Correct sequences were confirmed by Sanger sequencing followed by transformation of plasmids into ArcticExpress (DE3) competent cells. Xl-HMCES-3flag and xl-HMCES(E129A)-3flag proteins were expressed and purified as described above for xl-HMCES and xl-HMCES(E129A).

#### Generation of HMCES-DPCs

Crosslinking reactions with different HMCES variants were carried out in 10 μL reactions containing 8.02 μL reaction buffer (20 mM HEPES/KOH pH 7.5, 50 mM KCl, 10 mM MgCl_2_, 2 mM TCEP, 0.1 mg/mL BSA), 0.5 μL HMCES^FL^/HMCES^SRAP^ (prediluted to 40 μM in purification buffer D), 1 μL Cy5-labelled forward oligonucleotide (prediluted to 10 μM in DPC dilution buffer – 50 mM HEPES/KOH pH 7.5, 100 mM KCl, 10% glycerol, 0.4 mg/mL BSA) and 0.48 μL UDG (New England BioLabs), adding up to final concentrations of 2 μM HMCES^FL^/HMCES^SRAP^, 1 μM DNA and 0.1 U/μL UDG. Reactions were incubated for 1 h at 37°C. Crosslinking reactions with xl-HMCES (Figure S3B and S3C) were carried out identically as described above, except that the 10 μL reactions contained 1.5 μL xl-HMCES (prediluted to 13.2 μM in purification buffer D) and 7.02 μL reaction buffer. As standard and if not stated otherwise, a 30mer oligonucleotide containing a central dU (5’-Cy5-CCC AAA AAA AAA AAdU AAA AAA AAA AAA CCC-3’) was used for crosslinking. For generation of different DNA structures, 1 μL of corresponding reverse oligonucleotides (diluted to 12 μM in nuclease-free H2O) were added to the crosslinking reaction and incubated for 2 min at 37°C before the temperature was decreased by 1°C/min until 20°C was reached to allow annealing of the reverse oligo. For ssDNA samples a non-complementary reverse oligo was added (5’-AAA CCC CCC CCC CCA CCC CCC CCC AAA-3’), for ssDNA-dsDNA junction samples a 15mer reverse oligo was added (5’-GGG TTT TTT TTT TTT-3’), and for dsDNA samples a 30mer reverse oligo was added (5’-GGG TTT TTT TTT TTT ATT TTT TTT TTT GGG-3’). Reverse oligonucleotides were annealed prior to crosslinking for experiments shown in Figure 2A, S3A and S3B and after crosslinking for experiments shown in Figure 2C, 3B, 4A, 4B S2C, and S3C.

#### HMCES-DPC formation assays

For the experiments shown Figure S1B, the indicated HMCES variants were pre-diluted to 64, 32, 16, 8, 4, 2, 1, and 0.5 μM in purification buffer D prior to crosslinking. 0.5 μL of the pre-dilutions were added to the crosslinking reactions as described above resulting in final HMCES concentrations of 3.2, 1.6, 0.8, 0.4, 0.2, 0.1, 0.05, 0.025 μM. Otherwise, crosslinking reactions were carried out as described above. Reactions were stopped by addition of 5.5 μL LDS sample buffer and boiling for 1 min at 95°C. Samples were resolved on 4-12% SDS-PAGE gels. Gels were photographed using a BioRad Chemidoc MP system using appropriate filter settings for Cy5 fluorescence. Crosslinking was quantified using ImageJ by measuring the relative fraction of Cy5 signal in the DPC band.

For the experiments shown in Figure 2A and S3B, crosslinking reactions were set up as described above. Incubation and annealing of reverse oligonucleotides were performed in the absence of HMCES to generate desired DNA structures (ssDNA, ssDNA-dsDNA junction, dsDNA) before incubation with HMCES. Following annealing, 0.5 μL HMCES (prediluted to 40 μM in purification buffer D) was added to the reactions for Figure 2A or 1.5 μL xl-HMCES (prediluted to 13.2 μM in purification buffer D) for Figure S3B. Reactions were incubated for 0, 1, 5, 10, 20, or 30 min at 37°C before being stopped by addition of 5.5 μL LDS sample buffer and boiling for 1 min at 95°C. Samples were frozen in liquid nitrogen and stored at −80°C. Before resolving samples on 4-12% SDS-PAGE gels, samples were boiled again at 95°C for 30 s. Gels were photographed using a BioRad Chemidoc MP system using appropriate filter settings for Cy5 fluorescence. Quantification was performed using ImageJ by measuring the relative fraction of Cy5 fluorescence in the HMCES-DPC band.

#### HMCES-DPC release assays

For the experiments shown in Figure 1E and 1G, indicated HMCES variants were crosslinked to a 30mer oligonucleotide containing a central dU or dT (5’-Cy5-CCC AAA AAA AAA AAdU/dT AAA AAA AAA AAA CCC-3’) as described above. In parallel crosslinking reactions containing a 6-FAM-labelled 30mer oligonucleotide also containing a central dU (5’-6-FAM-CCC AAA AAA AAA AAdU AAA AAA AAA AAA CCC-3’) with 0.5 μL purification buffer D instead of protein were prepared and incubated at 37°C for 1 h as well. To inactive non-crosslinked HMCES, reactions containing the Cy5-oligonucleotide were incubated for 5 min at 60°C. In the following step, reactions containing the Cy5-oligonucleotide and HMCES were mixed 1:1 with reactions containing the 6-FAM-labelled oligonucleotide and incubated for 2 h at 37°C. To stop reactions, 11 μL of LDS sample buffer were added and reactions were boiled for 1 min at 95°C before analysis on 4-12% SDS-PAGE gels. Gels were photographed using a BioRad Chemidoc MP system using appropriate filter settings for Cy5 and 6-FAM fluorescence. Brightness and contrast were globally adjusted using ImageJ. 6-FAM- and Cy5-DPC formation was quantified using ImageJ.

For experiments shown in Figure 2C and S3C, indicated HMCES variants were crosslinked to a ssDNA oligonucleotide, as described above. Afterwards, corresponding reverse oligonucleotides were annealed as described above. Following annealing, 1 μL of the crosslinking reaction was added to 9 μL of master mix, resulting in a final buffer composition of 17.1 mM HEPES, 85.6 mM KCl, 3.1% glycerol, 5.5 mM TCEP, 2 mM MgCl_2_, and 0.1 mg/mL BSA. Reactions were either stopped directly after annealing (0 h), or after 1,2, 4, or 18 h incubation at 37°C by addition of 5.5 μL LDS sample buffer. Reactions were boiled for 1 min at 95°C before being frozen in liquid nitrogen and being stored at −80°C. Samples were boiled at 95°C for 30 s before being resolved on 4-12% SDS-PAGE gels. Gels were photographed using a BioRad Chemidoc MP system using appropriate filter settings for Cy5 fluorescence. Brightness and contrast were globally adjusted using ImageJ. Quantification was performed also using ImageJ by measuring the relative fraction of Cy5 fluorescence in the HMCES-DPC band.

For the experiments shown in Figure 3, indicated HMCES^SRAP^-variants were crosslinked as described above. Afterwards, corresponding reverse oligonucelotides were annealed to create a ssDNA-dsDNA junction, while a non-complimentary oligonucleotide was added in ssDNA conditions. HMCES^FL^ was pre-diluted to 20 μM in competition buffer (150 mM KCl, 50 mM HEPES, and 10% glycerol). The final assay was carried out in a reaction volume of 10 μL with 1 μL of the crosslinking reaction and 1 μL of pre-diluted HMCES^FL^, in a final buffer composition of 17.1 mM HEPES, 85.6 mM KCl, 3.1% glycerol, 5.5 mM TCEP, 2 mM MgCl_2_, and 0.1 mg/mL BSA. Reactions were incubated for 0, 1, 2, 3, 4, or 5 h at 37°C before being stopped by addition of 5.5 μL LDS sample buffer. The reactions were boiled for 1 min at 95°C before being frozen in liquid nitrogen and being stored at −80°C. After thawing samples were boiled at 95°C for 30 s before being resolved on 4-12% SDS-PAGE gels. Gels were photographed using a BioRad Chemidoc MP system using appropriate filter settings for Cy5 fluorescence. Brightness and contrast were globally adjusted using ImageJ. Quantification was done using ImageJ, by measuring the relative Cy5 signals of HMCES^FL^-DPCs and HMCES^SRAP^-DPCs.

#### APE1 incision assays

For the experiments shown in Figure 4, indicated HMCES^SRAP^-variants were crosslinked to a 30mer ssDNA as described above. Reverse oligonucleotides were annealed to generate a ssDNA-dsDNA junction or dsDNA. After annealing, 1 μL of the HMCES^SRAP^-DNA crosslinking reaction and 0.5 μL of APE1 (New England BioLabs) were added to 8.5 μL final reaction buffer, bringing final concentrations to 17.1 mM HEPES, 85.6 mM KCl, 3.1% glycerol, 5.5 mM TCEP, 2 mM MgCl_2_, and 0.1 mg/mL BSA. Reactions were either stopped directly (0 h), or after 1 or 18 h of incubation at 37°C by the addition of 5.5 μL LDS sample buffer. Reactions were boiled for 1 min at 95°C before being frozen in liquid nitrogen and being stored at −80°C. Samples were boiled again at 95°C for 30 s after thawing before being resolved on 4-12% SDS-PAGE gels. Gels were photographed using a BioRad Chemidoc MP system using appropriate filter settings for Cy5 fluorescence. Brightness and contrast were globally adjusted using ImageJ.

#### DNA binding assays

##### Electrophoretic mobility shift assays

HMCES^SRAP^-WT and variants were pre-diluted to 40, 10, and 2.5 μM in purification buffer D. Binding reactions were carried out in 10 μL with 0.5 μL of HMCES^SRAP^ dilutions, 1 μL of 1 μM Cy5-labelled 30mer dT-oligonucleotide and 8.5 μL reaction buffer (20 mM HEPES/KOH pH 7.5, 50 mM KCl, 10 mM MgCl_2_, 2 mM TCEP, 0.1 mg/mL BSA). Reactions were incubated for 20 min on ice before addition of 4 μL 6x Orange G loading dye. Samples were then resolved at 4°C on 6% native PAGE gels using 0.5x TBE as running buffer. Gels were photographed using a BioRad Chemidoc MP system using appropriate filter settings for Cy5 fluorescence. Brightness and contrast were globally adjusted using ImageJ.

##### Fluorescence polarization

HMCES^SRAP^-WT and variants were prediluted to 200 μM, 40 μM, 8 μM, 1.6 μM, 0.32 μM, 1.28 nM, and 0.256 nM in purification buffer D. Binding was carried out in 50 μL final volume with 5 μL of HMCES^SRAP^ pre-dilutions, 5 μL of 250 nM Cy5-labelled 30mer dT-oligonucleotide and 40 μL of reaction buffer (20 mM HEPES/KOH pH 7.5, 50 mM KCl, 10 mM MgCl_2_, 2 mM TCEP, 0.1 mg/mL BSA). Binding reactions were incubated for 20 min on ice before 10 μL of the reactions were pipetted into a 384-well microplate (Greiner Bio-One). Fluorescence polarization was measured using a Tecan Spark multimode microplate reader using appropriate filter settings for Cy5 fluorescence.

#### Preparation of oligonucleotide duplexes with AP-ICL

To generate the AP-ICL-containing oligonucleotide duplex, the complementary 5’-phosphorylated oligonucleotides (AP-ICL top: 5’-GCA CCT TCC GCT CdUT CTT TC-3’ and AP-ICL bottom: CCC TGA AAG AAG AGC GGA AG) heated for 5 min at 95°C in 30 mM HEPES-KOH pH 7.0, 100 mM NaCl, and annealed by cooling at 1°C per min to 18°C. The annealed duplex was treated with uracil glycosylase (NEB) in 1×UDG buffer for 2 h at 37°C followed by phenol/chloroform extraction and ethanol precipitation. The oligo duplex was suspended in 50 mM HEPES-KOH pH 7.0, 100 mM NaCl and then incubated at 37°C for 5 days to allow for cross-link formation. Cross-linked DNA duplex was separated on a 20% polyacrylamide, 1×Tris-borate-EDTA (TBE), 8 M urea gel and the cross-linked product was excised from the gel and eluted into TE (pH 8.0) buffer. Eluted DNA was concentrated by adding 4.5 times volume of 1-butanol, extracted with phenol:choloroform:isoamyl alcohol (25:24:1; pH 8.0), and precipitated with ethanol. The AP-ICL DNA oligo was then suspended in 10 mM Tris-HCl (pH 8.5) and stored at −80°C.

#### Preparation of cross-link DNA construct pICL-*lacO*A^P^

pICL-*lacO*^AP^ was prepared as described previously^22^. Briefly, the backbone plasmid (with 48 *lacO* repeats) was incubated with BbsI in NEBuffer 2.1 for 24 h at 37°C followed by phenol/chloroform extraction. The digested plasmid was further purified using a HiLoad 16/60 Superdex 200 column, which was equilibrated in TE pH 8.0 buffer. Fractions containing the linearized plasmid were pooled, precipitated in ethanol, and dissolved in 10 mM Tris-HCl pH 8.5. The abasic site interstrand cross-link (ICL^AP^)-containing duplexes were ligated into the linearized plasmid backbone using T4 DNA ligase (NEB). The ligated plasmid was dialyzed into TE pH 8.0 buffer and concentrated using an Amicon Ultra-15 10 kDa molecular weight cut-off filter unit. The covalently closed circular plasmids were futher extracted using the CsCl ethidium bromide method. Ethidium bromide was then removed from DNA by mixing with equal volume of saturated isobutanol. The purified pICL-*lacO*A^P^ was then dialyzed into TE pH 8.0, concentrated, snap frozen and stored at −80°C.

#### *Xenopus* egg extracts

The high-speed supernatant (HSS) and nucleoplasmic extracts (NPE) were prepared from *Xenopus laevis* eggs as described previously^22^. Briefly, six adult female *X. laevis* frogs were induced to produce eggs by injection with 500 IU hCG. Eggs were collected and dejellied in 1 L of 2.2% (w/v) cysteine, pH 7.7. Dejellied eggs were then washed with 2 L of 0.5× Marc’s Modified Ringer’s solution (2.5 mM HEPES-KOH, pH 7.8, 50 mM NaCl, 1 mM KCl, 0.25 mM MgSO4, 1.25 mM CaCl2, 0.05 mM EDTA) followed by 1 L of Egg Lysis Buffer (ELB) (10 mM HEPES-KOH, pH 7.7, 50 mM KCl, 2.5 mM MgCl_2_, 250 mM sucrose) supplemented with 1 mM DTT and 50 μg/mL cycloheximide. Eggs were then packed and crushed in the presence of 5 μg/mL aprotinin, 5 μg/mL leupeptin and 2.5 μg/mL cytochalasin B by centrifugation at 20,000 g for 20 min at 4°C. The soluble extract layer (the low-speed supernatant (LSS)) was collected and supplemented with 50 μg/mL cycloheximide, 1 mM DTT, 10 μg/mL aprotinin, 10 μg/L leupeptin and 5 μg/mL cytochalasin B. LSS was centrifuged in thin-walled ultracentrifuge tubes at 260,000 g for 90 min at 2°C in a TLS 55 rotor. Supernatant (HSS) was collected, aliquoted, snap frozen in liquid nitrogen, and stored at −80°C. To prepare NPE, LSS was prepared from eggs collected from 20 female *X. laevis* frogs as described above. LSS was then supplemented with 50 μg/mL cycloheximide, 1 mM DTT, 10 μg/mL aprotinin, 10 μg/mL leupeptin, 5 μg/mL cytochalasin B and 3.3 μg/mL nocodazole and centrifuged at 20,000 g for 15 min at 4°C. After removing lipids, the clarified cytoplasmic fraction was collected and supplemented with ATP regenerating mix (2 mM ATP, 20 mM phosphocreatine and 5 μg/mL phosphokinase) and 4,400 demembranated *X. laevis* sperm chromatin/μL to initiate nuclei formation. After ~90 min incubation at RT, reaction mixture was centrifuged for 3 min at 18,000 g at 4°C. The nuclei layer was collected from the top of the tubes and centrifuged at 260,000 g for 30 min at 2°C. The lipid layer was removed and the NPE fraction was collected, aliquoted, snap frozen in liquid nitrogen and stored at −80°C.

#### Antibodies and immunodepletion

Endogenous SPRTN protein was immunodepleted in HSS and NPE using anti-SPRTN antibodies (Rb31053) as previoulsy described^22^. Briefly, protein A Sepharose Fast Flow beads were washed thrice with PBS and then incubated with 3 volumes of SPRTN antibodies overnight at 4°C. The beads were then washed twice with PBS, once with ELB, twice with ELB supplemented with 500 mM NaCl, and thrice with ELB. HSS and NPE were immunodepleted by three rounds of incubation at 4°C for 60 min with protein A Sepharose-bound antibodies (5 volume extract per volume of beads). Extracts were centrifuged at 622 g for 30 s and the supernatants were collected.

#### Replication reactions

For replication reactions, licensing was conducted by incubating HSS with 7.5 ng/μL pICL-*lacO*^AP^ plasmid in the presence of 3 μg/mL nocodazole, 20 mM phosphocreatine, 2 mM ATP, 5 μg/mL creatine phosphokinase with or without [α-^32^P] dCTP for 30 min at RT. Replication was then initiated by mixing 2 volume of NPE mix (50% (v/v) NPE, 20 mM phosphocreatine, 2 mM ATP, 5 μg/mL creatine phosphokinase, 4 mM in ELB supplemented with 200 μM NMS-873 and 200 μM MG262) with 1 volume of licensing mix. Replication reactions were additionally supplemented with 250 nM WT xl-HMCES-3flag or 250 nM E129A xl-HMCES-3flag, as indicated. ^32^P-radiolabeled reactions were quenched by adding 1 μL of replication reaction to 6 μL of replication stop buffer (8 mM EDTA, 0.13% phosphoric acid, 10% ficoll, 5% SDS, 0.2% bromophenol blue, 80 mM Tris-HCl, pH 8.0) at the indicated time points followed by digestion with proteinase K (2.5 mg/mL) for 60 min at 37°C. Replication products were resolved on 0.8% agarose gel and visualized by phosphorimaging using a Typhoon FLA 9500 (GE Heathcare) and FujiFilm FLA 9500 user interface v.1.1. Images were analysed using ImageJ v1.53.

#### Plasmid pull-downs and immunoblotting

Plasmid pulldowns were performed as described previously^22^. Briefly, streptavidin-coupled magnetic Dynabeads (10 μL per pull down) were washed thrice with bead washing buffer (50 mM Tris-HCl pH 7.5, 150 mM NaCl, 1 mM EDTA pH 8.0, 0.02% Tween-20). Dynabeads were then incubated with biotinylated LacI (6 pmol per 10 μL of beads) at RT for 60 min. The beads were then washed thrice with pull-down buffer (20 mM Tris pH 7.5, 150 mM NaCl, 2 mM EDTA pH 8, 0.5% IPEGAL-CA630). 10 μL replication reaction was quenched into 400 μL of pull-down buffer and stored on ice. Samples were then incubated with 10 μL of LacI-coated streptavidin Dynabeads for at 4°C for 30 min on a rotating wheel. The beads were washed thrice with pull-down buffer followed by twice with Benzonase buffer (20 mM Tris pH 7.5, 20 mM NaCl, 2 mM MgCl_2_, 0.02% Tween-20). Beads were then suspended in 10 μL of Benzonase buffer containing 5 U Benzonase and incubated for 1 h at 37°C. 10 μL of 2×Laemmli loading buffer was added and the samples were incubated at 95 °C for 10 min. The supernatant was collected and resolved on a 10% Criterion TGX precast midi protein gel. HMCES-3flag protein was detected by immunoblotting with anti-flag antibody (1:10,000 dilution in PBST; Sigma, F1804-200UG) and rat anti-mouse secondary antibody peroxidase conjugate (1:20,000 dilution in PBST; Jackson ImmunoResearch Laboratories Inc., 415-035-166). SPRTN was detected using SPRTN anti-serum (Rb 31053; 1:5,000 dilution in PBST) and goat anti-rabbit secondary antibody peroxidase conjugate (1:20,000 dilution in PBST; Jackson ImmunoResearch Laboratories Inc., 111-035-003).

### QUANTIFICATION AND STATISTICAL ANALYSIS

Statistical details of each experiment (including the exact value of n, what n represents and precision measures) can be found in the figure legends.

